# The structure of an injectisome export gate demonstrates conservation of architecture in the core export gate between flagellar and virulence type three secretion systems

**DOI:** 10.1101/590273

**Authors:** Steven Johnson, Lucas Kuhlen, Justin C. Deme, Patrizia Abrusci, Susan M. Lea

**Author notes:** These authors made equal contributions to the work presented herein (Steven Johnson, Lucas Kuhlen). Structural Genomics Consortium, University of Oxford, Old Road Campus Research Build, Roosevelt Drive, Oxford OX3 7DQ, UK. Correspondence to: Susan M. Lea,.

## Abstract

Export of proteins through type three secretion systems (T3SS) is critical for motility and virulence of many major bacterial pathogens. Proteins are exported though a genetically defined export gate complex consisting of three proteins. We have recently shown at 4.2 Å that the flagellar complex of these three putative membrane proteins (FliPQR in flagellar systems, SctRST in virulence systems) assemble into an extra-membrane helical assembly that likely seeds correct assembly of the rod above. Here we present the structure of an equivalent complex from the more fragile Shigella virulence system at 3.5 Å by cryo-electron microscopy. This higher resolution structure reveals further detail and confirms the prediction of structural conservation in this core complex. Analysis of particle heterogeneity also reveals details of how the SctS/FliQ subunits sequentially assemble in the complex.

## Introduction

Virulence associated type three secretion systems (T3SS), also termed injectisomes, are bacterial nano-machines that facilitate the delivery of effector proteins directly into a eukaryotic host cell cytoplasm (Erhardt, Namba et al. 2010, Abrusci, McDowell et al. 2014). Injectisomes are closely related to the T3SS at the heart of the bacterial flagellum, and both of these classes are associated with the pathogenicity of a wide range of clinically relevant bacteria bacteria (Buttner 2012). T3SS vary significantly, being found in both Gram-negative and Gram-positive bacteria, and with further diversity defined by the existence of extracellular and periplasmic flagella. However, at the core of all T3SS is a basal body formed by circularly symmetric protein oligomers spanning the inner membrane, from which helical flagellar filaments or injectisome needle structures project (Macnab 2003, Erhardt, Namba et al. 2010, Deng, Marshall et al. 2017). Proteins associated with the cytoplasmic face of the basal body select proteins for export that are then transferred to a set of 5 membrane associated proteins located at the center of the inner-membrane ring. These components (FliP, FliQ, FliR, FlhB, FlhA in the flagellar system and SctR, SctS, SctT, SctU, SctV in injectisomes) are collectively termed the export apparatus (EA) and are absolutely required for the translocation of substrates across the bacterial envelope (Wagner, Konigsmaier et al. 2010, Fabiani, Renault et al. 2017, Fukumura, Makino et al. 2017).

We have recently demonstrated that a subset of these proteins assembles into a core export-gate complex, and reported the structure of a flagellar FliP_5_Q_4_R_1_ (hereafter termed FliPQR) complex at 4.2 Å structure by cryo-electron microscopy (Kuhlen, Abrusci et al. 2018). Strikingly placing the FliPQR structure into lower resolution basal body structures revealed that this complex, built from three putative membrane proteins and purified from membranes when expressed in isolation of the rest of the T3SS, physiologically exists in an extra-membrane location at the core of the basal body (Worrall, Hong et al. 2016, Kuhlen, Abrusci et al. 2018)}. This positioning of the complex, in conjunction with the observation that it exhibits helical symmetry, suggested that it seeds assembly of the axial helical components that culminate in the flagellum or needle. The high level of sequence conservation within all T3SS implied that this complex would be similarly assembled in virulence T3SS. Native mass spectrometry (nMS) of purified virulence system complexes supported this, revealing a core SctR_5_T_1_ complex equivalent to FliP_5_R_1_ (Kuhlen, Abrusci et al. 2018). However, the number of SctS subunits was highly variable in samples and seemed to reflect lower stability of these complexes c.f. the flagellar ones (Kuhlen, Abrusci et al. 2018). This decreased stability meant that determination of a structure from a non-flagellar export gate complex was not previously possible.

Here we present a re-refinement of the original FliPQR data to higher resolution (3.65 Å) and also the structure of the equivalent SctRST complex from a virulence T3SS (3.5 Å), revealing the high level of structural conservation within this core complex between flagellar and virulence T3SSs. The greater fragility of the virulence complex c.f. the flagellar complex means that our sample contains a variety of differently assembled complexes that differ in the number of SctS (FliQ) subunits associated with the complex, supporting a model for sequential assembly of the complex coupled with extrusion from the inner membrane.

## Results & Discussion

Re-refinement of the original *Salmonella enterica* serovar Typhimurium (*S.* Typhimurium) flagellar FliPQR data (Kuhlen, Abrusci et al. 2018) using RELION-3 (Zivanov, Nakane et al. 2018) yielded a significant improvement in the resolution of the density, as estimated from both FSC curves and the level of detail visible in the volume (Fig. 1). We have remodeled and re-refined the coordinates into the new volume and, although there are no substantial differences, the level of confidence in these coordinates as accurately representing the biological object is clearly increased.

**Figure 1.**
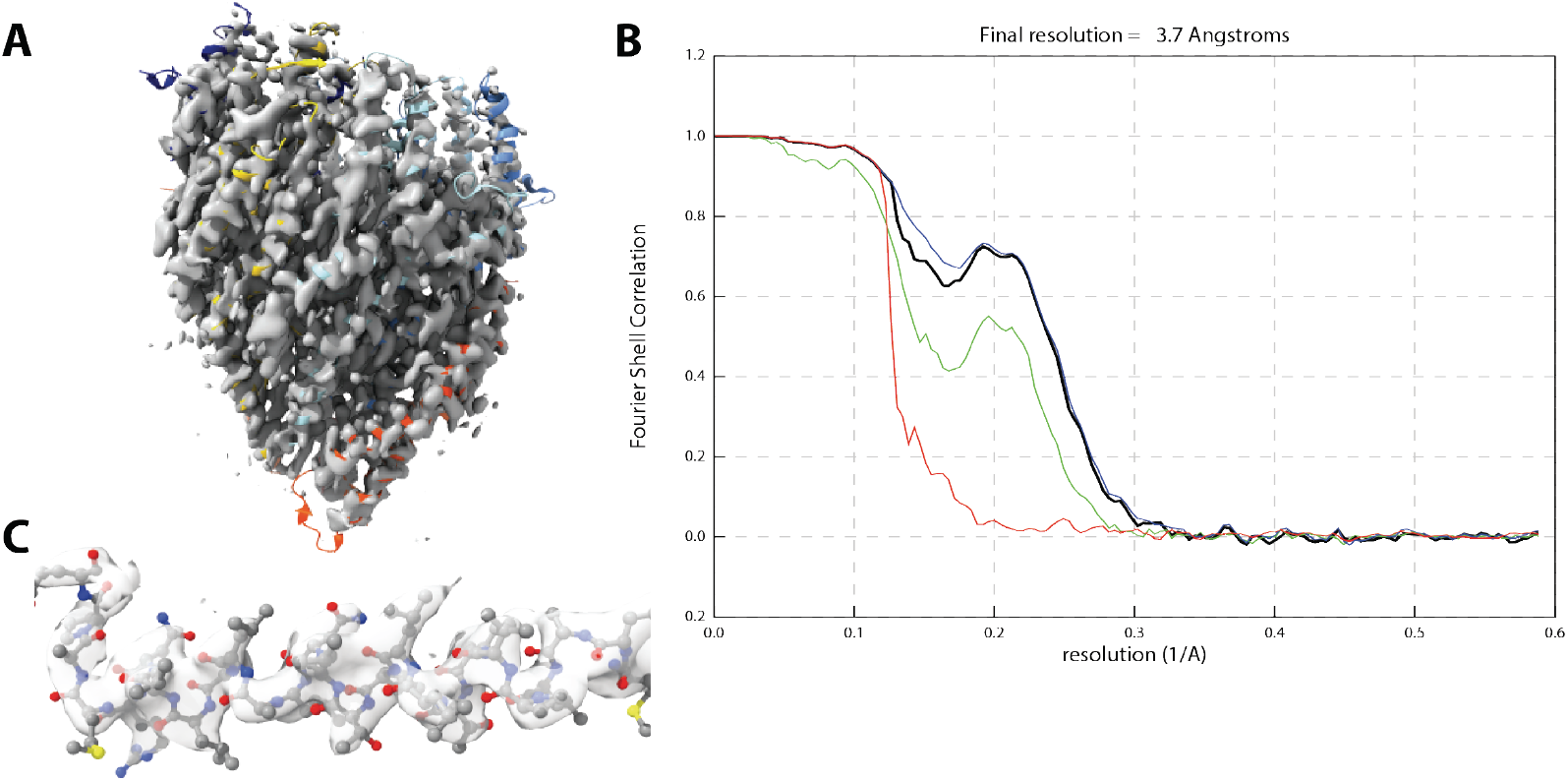
Improved resolution of Salmonella FliPQR reconstruction. (A) Volume obtained after re-refinement in RELION-3 with particle polishing and per-particle CTF refinement (3.65 Å, EMD-4733) (B) closeup of region 190-210, chain F (FliR) to illustrate quality of density (C) FSC curve black – FSC corrected, green – FSC unmasked maps, blue - FSC masked maps, red – FSC phase randomised

Following our determination of the structure of the flagellar export gate complex we expressed a variety of virulence export gate complexes using the same strategy of expression of the complete operon with a dual-strep tag on the C-terminus of the SctT (FliR) component. Many of the systems proved fragile, with little to no SctS (FliQ) associated with the purified complexes (Kuhlen, Abrusci et al. 2018), but the *Shigella flexneri* export gate (Spa24, Spa9, Spa29 – hereafter referred to as SctRST) could be purified at sufficient levels to allow structure determination by single particle cryoEM (Fig 2A). As previously proposed (Kuhlen, Abrusci et al. 2018) based on the sequence conservation in all three components (33% identity to the *S.* Typhimurium FliPQR across the operon), the structure conclusively demonstrates the structural conservation at the core of type three secretion systems (Fig. 2B, C).

**Figure 2.**
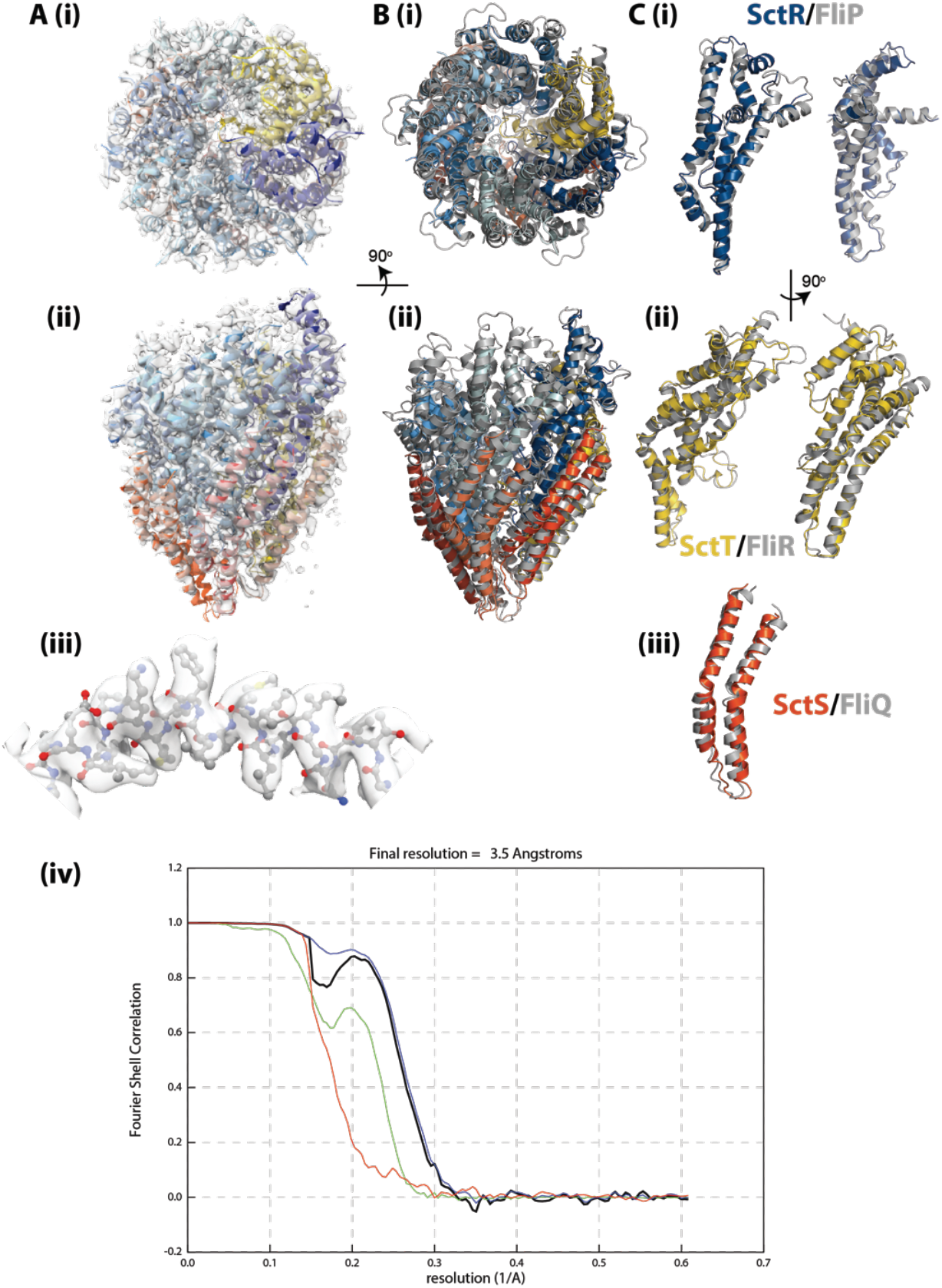
Structure of Shigella flexneri export gate (SctRST) and comparison with Salmonella FliPQR. (A) Different views of the coordinates within the 3.5 Å volume (EMD-4734) are shown (i) and (ii) show two views of the whole assembly related by a 90° rotation with the coordinates shown in a cartoon representation and coloured (as previously defined) with five copies of SctR in shades of blue, four copies of SctS in shades of red and with SctT in gold (iii) shows a close up of a region of SctR with all atoms shown (iv) FSC curve for reconstruction colors as in Fig. 1. (B) Overlay of the Shigella flexneri SctRST (coloured as in A) and Salmonella FliPQR (grey) with both shown in a cartoon representation. Views are as in A(i),(ii). (C) Individual overlays are shown for example of each type of chain extracted from the complex, coloured as in B.

As suggested by the nMS analysis (Kuhlen, Abrusci et al. 2018), the biggest difference between the flagellar and injectisome export gates was observed in the Q/S subunits. The highest resolution SctRST map was generated from ~ 200000 particles and displayed density for 4 copies of SctS, but with evidence that the 4th copy, corresponding to the lowest position in the helical assembly, was sub-stoichiometrically occupied. Further 3D classification of this particle set produced a series of lower resolution maps (Fig. 3A) highlighting the compositional heterogeneity within the SctRST sample, with evidence for SctR_5_S_2_T_1_, SctR_5_S_3_T_1_ and SctR_5_S_4_T_1_ complexes. Furthermore, the order in which the SctS copies were occupied clearly progressed down the helical assembly. This observation is in agreement with the nMS data and supports the hypothesis that the complex is assembled sequentially, with the SctS component being the last to be added.

**Figure 3.**
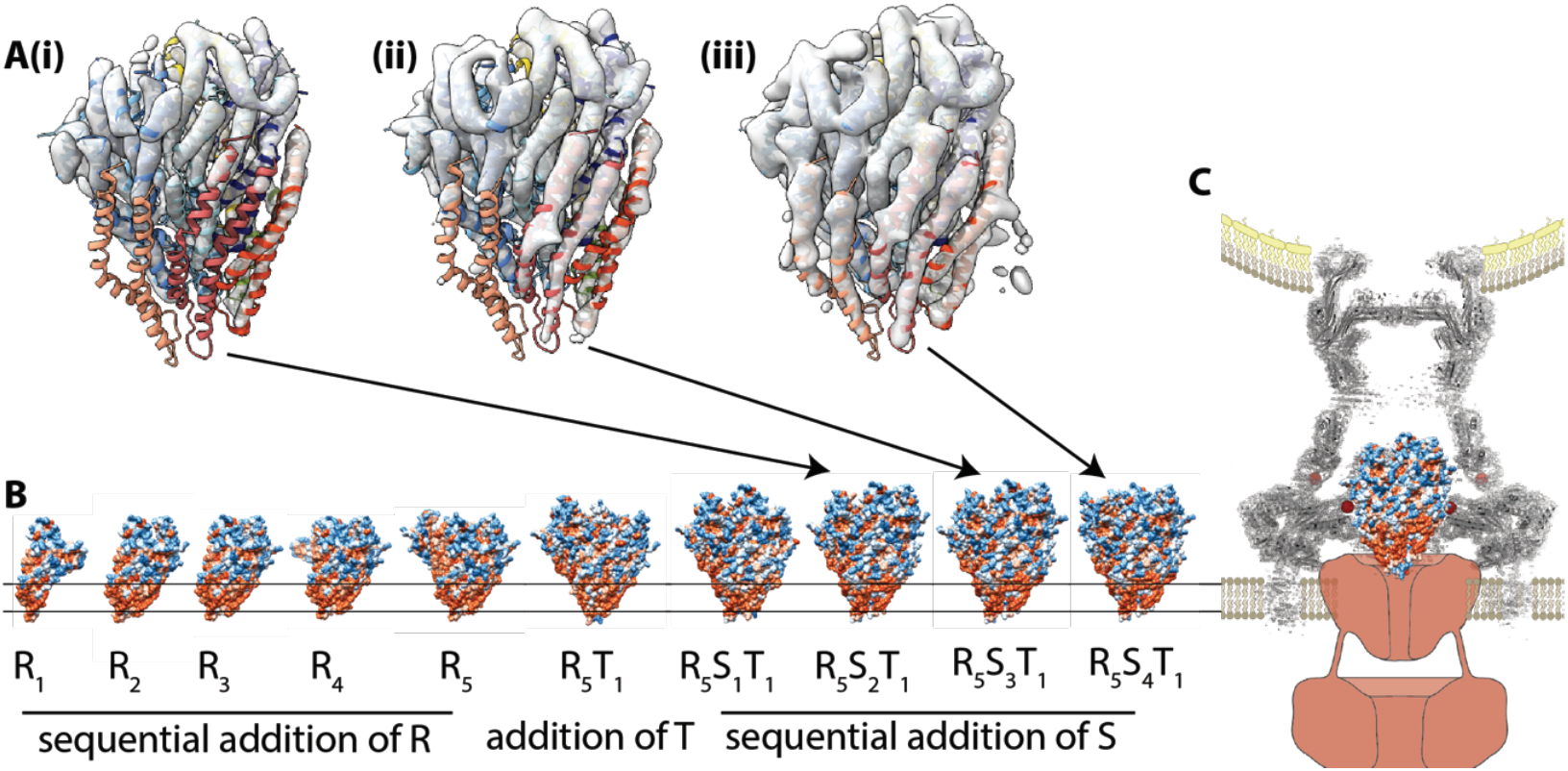
Model for assembly of the T3SS export gate (A) 3D classification of the SctRST particles leads to volumes with different numbers of SctS subunits attached (i) SctR_5_S_2_T_1_ (ii) SctR_5_S_3_T_1_ and (iii) the fully assembled SctR_5_S_4_T_1_. The Full assembly is shown as a cartoon trace within each volume with the absent SctS subunits seen to lack density at the contour level sufficient to cover the subunits present. (B) The height of the surface-exposed hydrophobic patches (orange) on the sub-complexes suggests sequential assembly in the order shown with the complex pushing out of the inner membrane into the periplasmic space (C) once the remaining T3SS basal body components assemble the export gate is found above the inner membrane at the core of the basal body (grey cartoon & density Worrall et al. 2016) with the SctV/FlhA component (red cartoon) assumed to form the channel in the inner membrane.

To attempt to further understand how this ultimately extra-membranous complex is assembled in the membrane we analysed the hydrophobic surfaces revealed when subunits are sequentially removed (Fig. 3B) from the complex. This analysis combined with the knowledge that the complex needs to be positioned ready for extraction from the membrane to allow its incorporation into the full T3SS basal body (Fig. 3C), suggests an ordered assembly process. This would begin with sequential assembly of five copies of SctR, followed by completion of a closed structure by addition of the single copy of SctT. Creation of the R/T interface would trigger sequential addition of the four copies of SctS. This raises interesting questions of how the operon order relates to the protein complex assembly, as this implies that the order of protein assembly does not match the order of the genes within the operon.

## Materials & Methods

### Materials

Chemicals were obtained from Sigma-Aldrich unless otherwise specified. The detergent lauryl maltose neopentyl (LMNG) was obtained from Anatrace.

### Recombinant protein expression

*Shigella flexneri* SctRST (Spa24/Spa9/Spa29) was recombinantly expressed in *Escherichia coli* and extracted and purified in LMNG as described previously (Kuhlen, Abrusci et al. 2018). Briefly, BL21 cells transformed with plasmid pT12_Spa24929 were grown overnight in terrific broth supplemented with kanamycin (60 ug/ml) and rhamnose monohydrate (0.1%) at 37 °C. The cells were spun down, lysed and the membranes pelleted by ultracentrifugation. The membranes were dissolved in 1% LMNG and the protein was purified by StrepTrap and size exclusion chromatography.

### CryoEM Grid preparation and Single Particle Data Collection

Purified SctRST in 0.01% LMNG in TBS (100 mM Tris, 150 mM NaCl, 1 mM EDTA, pH 8) was concentrated to 16.8 mg/ml and diluted to 8.4 mg/ml. 3 ul of sample were applied to glow-discharged holey carbon-coated grids (Quantifoil 300 mesh, Au R1.2/1.3), adsorbed for 5 s, blotted for 3 s at 100% humidity at 22°C and frozen in liquid ethane using a Vitrobot Mark IV (Thermofisher). All EM data were collected using a Titan Krios (Thermofisher) operating at 300kV. 3741 movies were collected on a K2 Summit detector (Gatan) in counting mode using a sampling of 0.822 Å/pixel (calibrated by prior determination of the structure of apo-ferrritin), with 2.4 e^-^Å^-2^frame^-1^ over 20 frames.

### Structure Solution

All steps in the structure solution were carried out using RELION-3 (Zivanov, Nakane et al. 2018) unless otherwise stated. Motion correction of the movies was carried out using MotionCor2 (Zheng, Palovcak et al. 2017), as implemented within RELION-3(Zivanov, Nakane et al. 2018), with dose weighting. CTF estimation was carried out using CTFFIND4 (Rohou and Grigorieff 2015). 775073 particles were extracted in RELION-3 using boxes picked using SIMPLE (Reboul, Eager et al. 2017). Reference free 2D classification was used to select 477653 good particles. 3D classification with 4 classes was then carried out using the *S.* Typhimurium FliPQR structure (Kuhlen, Abrusci et al. 2018) low pass filtered to 40 Å as a reference. The best class (212561 particles) was then used in 3D auto-refinement, followed by Bayesian polishing and CTF parameter refinement. A final gold-standard refinement produced the final map with a resolution of 3.5 Å after PostProcess masking and B-factor sharpening. Model building was initially carried out using CCP4-Buccaneer, followed by manual building in Coot (Brown, Long et al. 2015). The structure was refined using phenix.real_space_refine (Afonine, Poon et al. 2018).

**Table 1.**
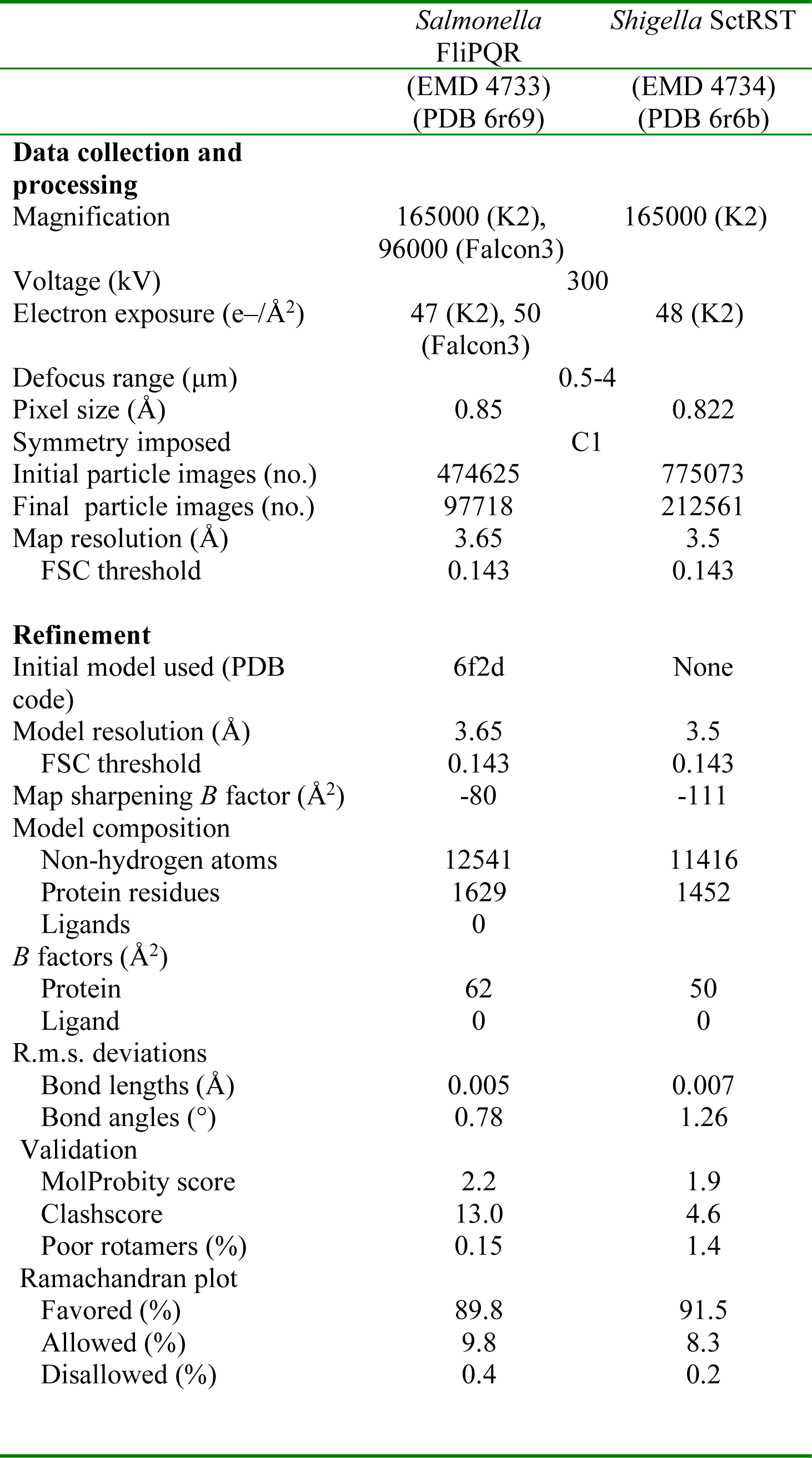
Cryo-EM Data Collection, refinement and validation statistics

## Database Deposition

Volumes and coordinates have been deposited in the EMDB and PDB respectively. *Salmonella* FliPQR EMD-4733 and PWD-ID 6R69, *Shigella* SctRST EMD-4734, PDB 6r6b

## Acknowledgements

We thank E. Johnson & A. Costin of the Central Oxford Structural Microscopy and Imaging Centre for assistance with data collection. H. Elmlund (Monash) is thanked for assistance with access to SIMPLE code ahead of release. The Central Oxford Structural Microscopy and Imaging Centre is supported by the Wellcome Trust (201536), The EPA Cephalosporin Turst, the Wolfson Foundation and a Royal Society/Wolfson Foundation Laboratory Refurbishment Grant (WL160052). Work performed in the lab of S. M. Lea was supported by a Wellcome Trust Investigator Award (100298) and an MRC programme grant (M011984). LK is a Wellcome Trust PhD student (109136)

## Author Contributions

S.J.:.Designed, supervised and performed experiments. Characterisation of protein complexes. CryoEM data analysis, structure determination and analysis. Wrote manuscript with SML.

L.K.: Performed experiments. Strain and plasmid construction, complex purification, native mass spectrometry, cryoEM grid optimisation, cryoEM data analysis and model building and analysis.

P.A.: Performed experiments. Strain and plasmid construction

J.D.: Performed experiments. CryoEM grid optimisation and data collection

S.M.L.: Designed, supervised and performed experiments. CryoEM data optimisation and collection, data and structure analysis. Wrote paper and prepared figures with SJ.

